# Molecular mechanisms regulating the pH-dependent pr/E interaction in yellow fever virus

**DOI:** 10.1101/2022.12.06.519383

**Authors:** E. Crampon, E. Covernton, M.C. Vaney, M. Dellarole, A. Sharma, A. Haouz, P. England, J. Lepault, S. Duquerroy, F.A. Rey, G. Barba-Spaeth

## Abstract

Flavivirus particles bud in the ER of infected cells as immature virions composed of 180 heterodimers of glycoproteins prM and E, associated as 60 (prM/E)_3_ trimeric spikes. Exposure to the mildly acidic pH of the TGN results in dissociation of the trimeric spikes followed by reassociation of the prM/E protomers into 90 dimers organized in a characteristic herringbone pattern. The furin site in prM is exposed in the dimers for maturation of prM into M and pr. For flaviviruses such as the tick-borne encephalitis virus (TBEV) as well as for dengue virus, it was shown that at neutral pH pr loses affinity for E, such that it dissociates from the mature particle as soon as it reaches the external milieu, which is at neutral pH. Using a soluble recombinant form of E (sE) and pr from yellow fever virus (YFV), we show here that the affinity of pr for recombinant E protein remains high even at neutral pH. The X-ray structure of YFV pr/sE shows more extensive inter-chain hydrogen bonding than does the dengue or TBEV, and also that it retains the charge complementarity between the interacting surfaces of the two proteins even at neutral pH. We further show that pr blocks sE flotation with liposomes when exposed at low pH at a 1:1 stoichiometry, yet in the context of the virus particle, an excess of 10:1 pr:E ratio is required to block virus/liposome fusion. In aggregate, our results show that the paradigm obtained from earlier studies of other flaviviruses does not apply to yellow fever virus, the flavivirus type species. A mechanism that does not rely solely in a change in the environmental pH is thus required for the release of pr from the mature particles upon release from infected cells. These results open up new avenues to understand the activation mechanism that yields mature, infectious YFV particles.

## INTRODUCTION

Enveloped viruses use membrane fusion protein (MFP) to mediate viral fusion with the host cell. The majority of MFPs belong to three structural classes, I, II, or III. Flaviviruses have class II MFPs carrying an elongated ectodomain divided into three distinct □-sheet rich domains (DI, DII, DIII), a stem region, and are anchored to the viral membrane by C-terminal trans-membrane (TM) domains (1). Their folding in the ER of the infected cell is assisted by an accompanying protein (AP) which acts as a chaperone. The MFP/AP heterodimer is the building block at the surface of the mature virus, with the AP positioned to protect the fusion loop (FL) of MFP, the hydrophobic region responsible for the insertion into the host membrane. Flaviviruses are the only exception. Their viral particle is indeed constituted by homodimers of the MFP envelope (E) protein tightly organized in a herringbone pattern with the FL buried at the homodimer interface. The pre-membrane (prM) protein is the flaviviruses AP protein. It gets cleaved by furin during flavivirus maturation in the trans-Golgi network (TGN) into M, that remains anchored by its TM domains to the viral membrane underneath the E homodimer, and pr moiety that interacts with the FL of E to prevent premature triggering of viral fusion in the acidic environment of the TGN (2), (3). The necessity to protect the FL and, at the same time, to have a particle ready to fuse after receptor-mediated endocytosis, has pushed the flaviviruses to evolve concerted strategies based on conformational changes of the E/prM complex driven by a low pH-triggering switch (4). During viral entry, the acidic pH of the endosome triggers a dimer-to-trimer transition of the E protein resulting in an exposure of the fusion loop at the tip of the trimer for insertion into the host membrane and successive viral fusion. During virus secretion in the secretory pathway, a trimer-to-dimer transition brings the trimer of E/prM heterodimers, that form as the noninfectious immature virus buds in the ER, to an E dimer with M underneath and pr on top, associated to the fusion loop. In the TGN the furin protease cleaves pr-M but pr remains associated to the pre-mature particle. When the viral particle is released in the neutral pH extracellular environment, pr is then removed from the virus which is now infectious and ready to begin a new cycle (5),(6).

The mechanism regulating these transitions is not fully understood but a recent work from Vaney and coll. showed how for TBE, during the transit across the secretory pathway, the 150 loop and the N-terminal of the E protein act in coordination with the pr protein to assure protection of the FL in the transition from low to neutral pH. It is indeed the movement of the 150 loop towards the N-terminal of E at neutral pH that actively expels pr from its binding site (7). These regions show structural conservation between flaviviruses suggesting a common mechanism of action, however, the determinants of their interaction may vary (i.e. the length of the 150 loop or the presence of glycosylation) and may result in differences in infectivity and/or pathogenicity (8).

Our work describes the interaction of pr/E for yellow fever virus (YFV) and identifies a unique interaction of pr/E at neutral pH, absent in the other flaviviruses. We show the structural basis of this interaction, relying on an extensive inter-chain hydrogen bonds with interactions specific to YFV. At low pH pr prevents E insertion into membranes and blocks viral fusion as for the other flaviviruses. However, at neutral pH, the pr/E interactions are weakened but still present suggesting the necessity of additional mechanisms for the release of pr from the mature particle.

## RESULTS

### Interactions of YFV pr and E proteins

To produce correctly folded YFV sE protein, we used the same strategy that we previously adopted for the production of dengue sE protein expressing the prME region as it is in the viral polyprotein (9). This type of construct assures the secretion of soluble E (sE) while M remains membrane-anchored in the cell and pr, cleaved by furin in the TGN, dissociates from sE when the complex reaches the extracellular milieu. Similar constructs for dengue and tick-borne encephalitis (TBE) viruses resulted indeed in secretion of the soluble sE protein (9), (10). In the case of YFV instead, we obtained a stable pr/sE complex even in the absence of covalent linker, which indeed was necessary for the production of DENV2 pr/sE complex crystallized previously (11). We obtained crystals for the YFV wild type Asibi pr/sE complex that yielded a structure to 2.7Å resolution and refined to free R factors of 19% (Fig. 1A and Suppl. Table S1). This structure shows the YFV pr/sE complex as it is supposed to be in the secretory pathway after furin cleavage.

**Figure 1:**
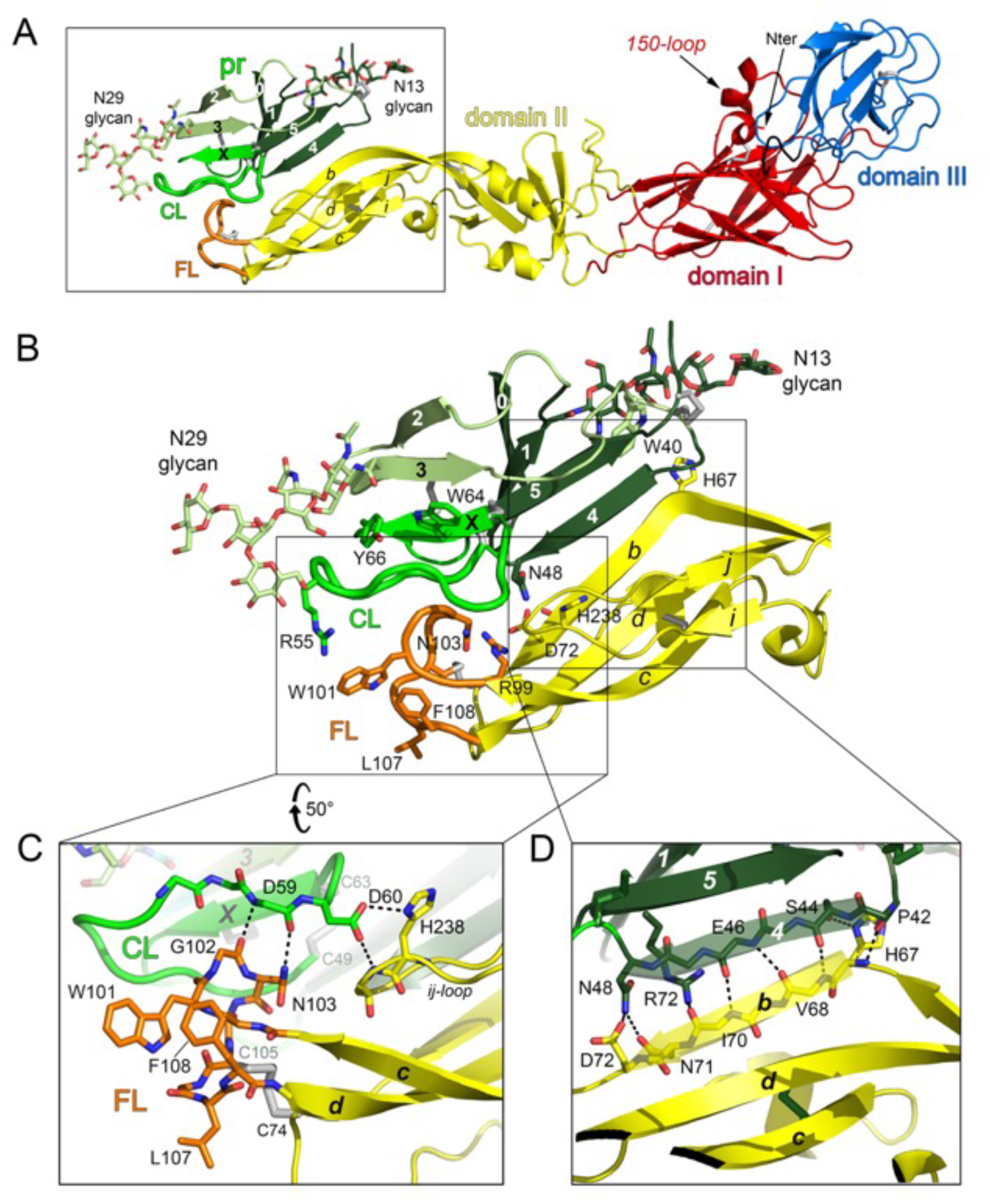
Structure of the pr/sE complex. **(A) Structure of the pr/sE complex**. E is colored according to the classical flavivirus E domains DI, Dll and Dill in red, yellow and blue respectively. The fusion loop (FL) is in orange, pr is in green and is glycosylated in positions Asn13 and Asn29 as indicated. The black arrows point the 150-loop and the N-terminal residue. **(B) The framed region in (A) is shown**. Residues of the fusion loop (FL) are displayed as sticks. E β-strands of domain II are labeled. The pr molecule is displayed with three shades of green, with the two beta-sheets sandwich colored dark and light green and with the capping-loop (CL) displayed thicker in bright green, pr is glycosylated in positions Asn13 and Asn29 as indicated. Residues discussed in the text are labelled and displayed as sticks and atom color-coded. In brief, Trp40 packs against Asn13 while Tyr66 and Arg55 pack against Asn29, stabilizing CL. **(C-D) The framed regions in (B) show close-views of the pr/sE interactions**. The residues that interact between pr and E are displayed as sticks and atom color-coded. **(C)** Interactions of residues of cd-loop (FL, in orange) and *ij-* loop (in yellow) with the residues of the pr-CL (in green). The residues that interact between pr and E are displayed as sticks and atom color-coded. The residues E-His238 and pr-Asp60 are strictly conserved among all the flaviviruses. **(D)** Hydrogen bonded network between the *β*-strand β4 of pr (in dark green) and the *β*-strand *b* of E (in yellow) involving main-chain and side-chain residues. The residues are labelled, and the directions of the strands are shown in transparency.

Although YFV E is not glycosylated, to sites of glycosylation, Asn13 and Asn29, are presentin YFV pr (Fig. 1A,B). Asn13 glycan, a specific glycosylation site in YFV group, packs against Trp40. The glycan Asn29 glycan is located on *β*-strand *β*3 in a location spatially just nearby just the DENV pr glycosylation Asn69 (YFV-Tyr66, *β*-strand *β*x). In YFV the glycan packs against Tyr66 and Arg55 stabilizing the capping loop conformation (CL, shown in bright green in Fig. 1B). The capping loop is a protruding loop that wraps around the sE fusion loop (FL, shown in orange in Fig.1). Its conformation is stabilized by a disulfide bridge between Cys49 and Cys63, and hydrophobic packing interactions with Trp64, Tyr66 and the Asn29 glycan chain (Fig. 1B). pr-CL makes multiple polar and hydrophobic contacts including many main-chain/main-chain interactions, some of them are conserved in DENV2 (PDB 3C5X) and TBEV (PDB 7QRE) pr/sE structures (Table 1). In particular, residue pr-Asp60 makes, among all the flaviviruses, a strictly conserved salt-bridge interaction with sE-His238 stabilizing the *ij*-loop of E-domain II (Fig. 1C). This interaction was previously described in the pr/sE complex of DENV2 (PDB code 3C5X) (11) and TBEV (PDB 7QRE, 7QRF) (7). In addition, pr and sE interaction is further stabilized by main-chain interactions and side-chains H-bonds between pr-Ser44 and sE-His67 or pr-Asn48 and sE-Asn71/Asp72 at both ends of the *β*-strands (Fig. 1D).

**Table 1.**
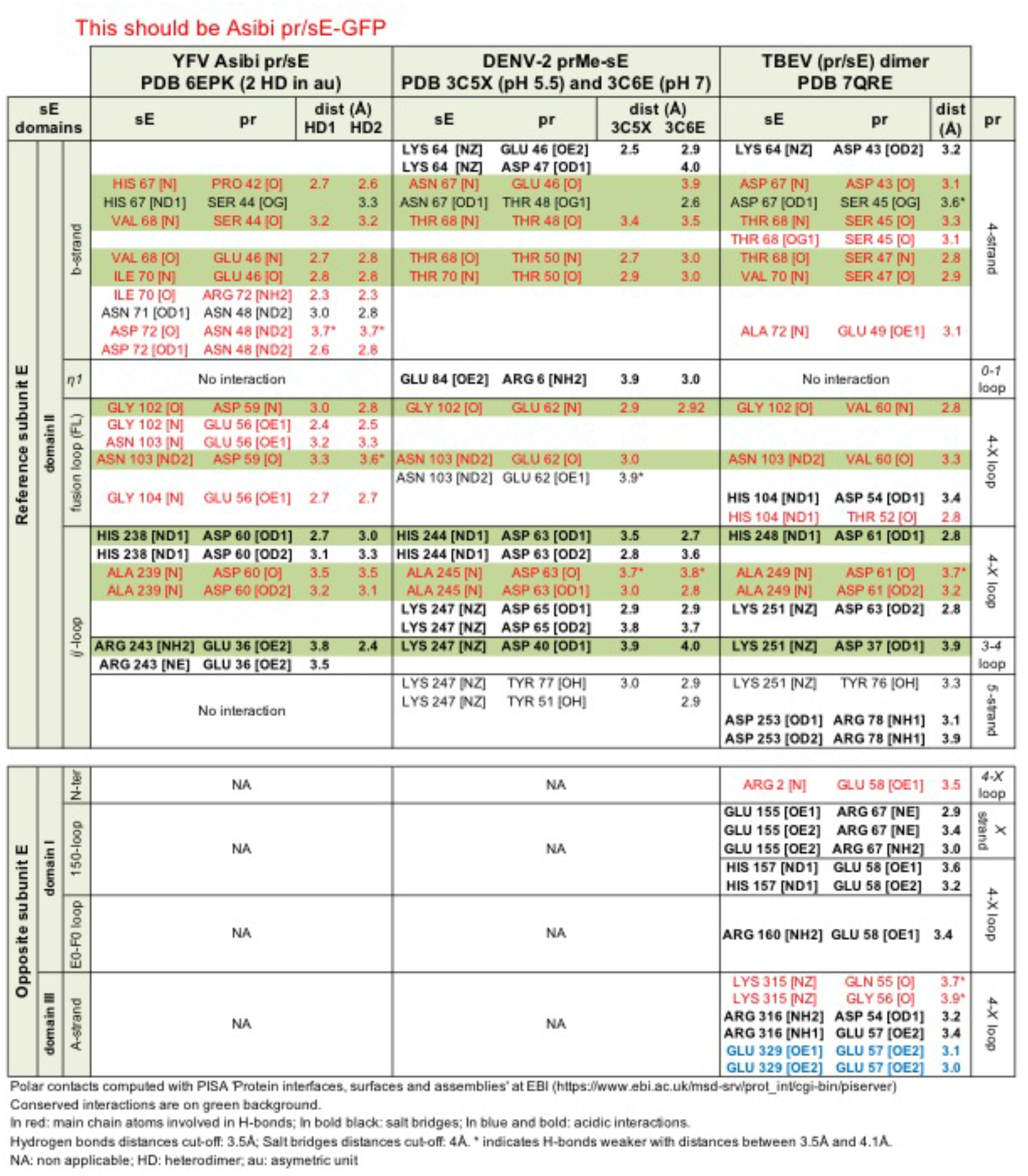
Comparison of polar pr/sE interactions in YFV - DENV - TBEV.

Thus, the tight association of YFV pr/E complex is supported by conserved interactions, also present in DENV2 and TBEV (highlighted by the green background in Table 1 and interactions specific to YFV involving the Ile70-Asp72 region (Fig. 1D and Table 1).

### Chaperone role of pr and YFV pr/sE interactions

To perform functional studies on YFV soluble E protein (sE) and its interactions with membranes, we had to separate the pr/E complex. A first attempt was done using anion exchange chromatography with a NaCl gradient. We were able to separate two peaks, one still containing the pr/sE complex and the other one containing sE alone (sE’) (Suppl. Fig. S1). However, further functional analysis of the protein sE’ eluting at 400mM NaCl, revealed that this protein was unable to insert into liposomes at acidic pH (see experimental details below) and it was probably a misfolded form of sE. We then used 8M urea for denaturation of the pr/sE complex eluting at 240mM NaCl, followed by renaturation with extensive dialysis against Tris 20mM pH 8.0 of the separated E and pr proteins. To simplify the protein preparation for the functional studies, we also decided to express in S2 cells the pr protein alone and the sE protein without prM. The yields of sE produced in absence of prM were sensibly lower and the SEC profile showed a large heterogeneity of the produced protein (Fig. 2A and B) confirming the chaperone role of prM for YFV E protein. Comparison with the SEC profile of the protein produced by the prME construct identified peak 3 as the corrected folded protein (Fig. 2A). Both peak 3 and sE obtained after urea treatment of pr/sE complex were used for further functional studies.

**Figure 2:**
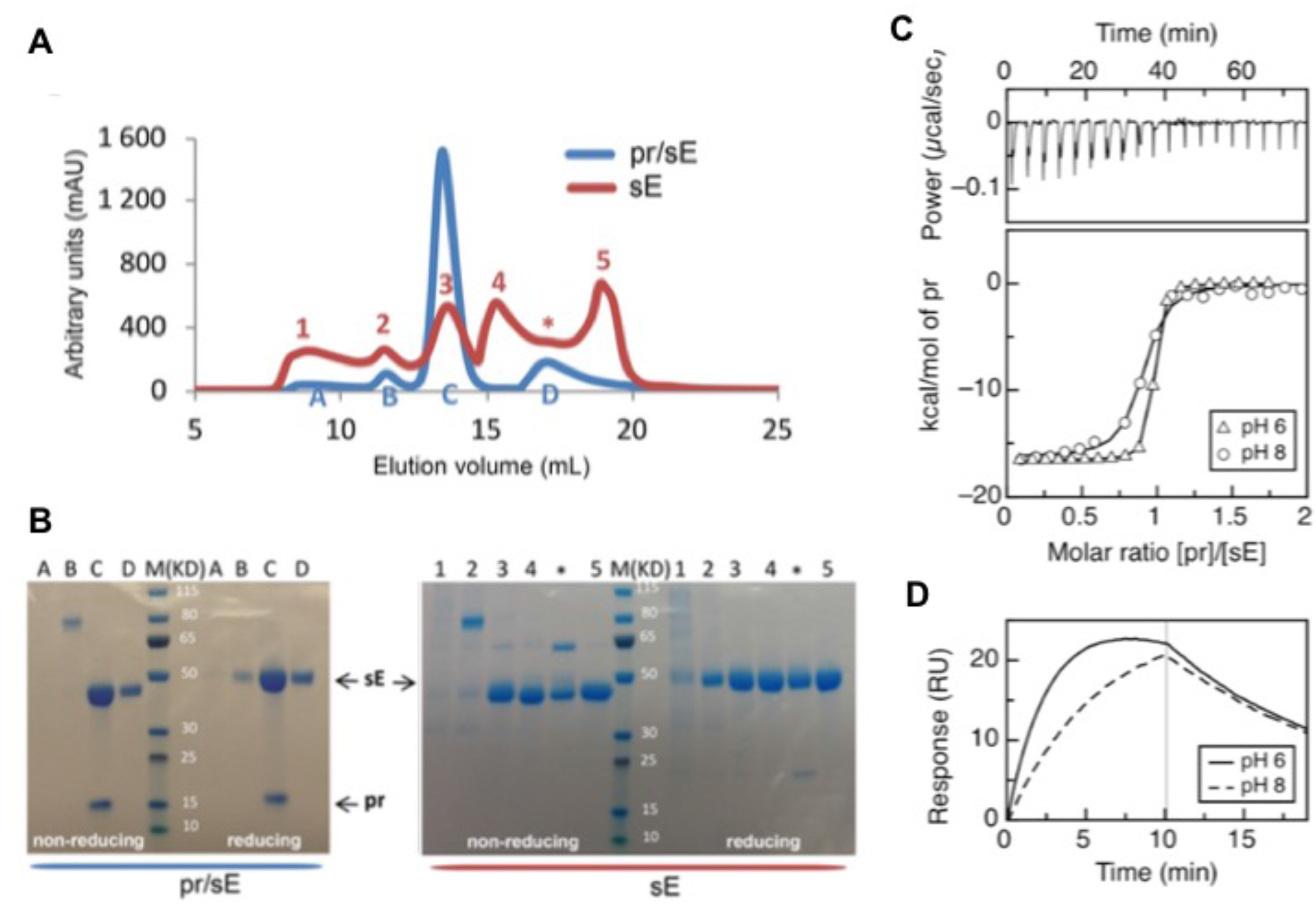
Chaperone activity of pr on the E protein and biophysical characterizations of the pr/sE interaction. **(A)** SEC profiles of YFV sE expressed with prM (in blue), showing 4 peaks, labelled A, B, C and D, and of sE expressed without prM (in red), showing 6 peaks labelled 1, 2, 3, 4, * and 5. Proteins were produced in S2 cells and first purified by affinity chromatography and then analyzed by size exclusion chromatography (SEC). **(B)** SDS-PAGE analysis of the protein present in each peak under reducing and non-reducing conditions, as indicated. Aliquots from each peak were run under non-reducing (without DTT) or reducing (+DTT) in an SDS-PAGE and stained with Coomassie blue. The asterisk in the profile of sE indicates a shoulder of peak 4 that was treated separately. Molecular masses of marker proteins are listed in kilodaltons. **(C) Isothermal titration calorimetry**. sE:pr binding isotherms (bottom panel), recorded at pH 6.0 and pH 8.0, shown as triangles and circles, respectively, resulting from integration of the specific heats with respect to time as shown for pH 8.0, top panel. **(D) Surface plasmon resonance**. Example of sE:pr association and dissociation kinetics corresponding to injections of pr at 12.5 nM over immobilized sE, respectively at pH 6.0 (solid line) and pH 8.0 (dashed line).

Differently to previous studies with dengue virus, which had shown very weak or no pr/sE interactions at neutral pH, we observed a stable pr/sE complex at pH 8.0. We measured the affinity of YFV pr/sE interaction at pHs 6.0 and 8.0 by two different methods, isothermal calorimetry (ICT, Fig. 2C) (12) and surface plasmon resonance (SPR, Fig. 2D). The dissociation constants (K_D_) obtained by the two independent methods followed a similar trend: a K_D_ under 10 nM at pH 6.0 (8.5nM by ITC and 6.2nM by SPR) and about five times higher at pH 8.0 (58.8nM by ITC and 19.7nM by SPR) (Table 2), indicating a pH sensitive interaction. In previous studies of the interaction between DENV2 pr and sE, although a K_D_ had not been reported, SPR experiments revealed undetectable or no binding at pH 8.0 (13), whereas in the case of YFV we find an affinity still under 100 nM under these pH conditions, indicating a real difference in the two viruses.

**Table 2.**
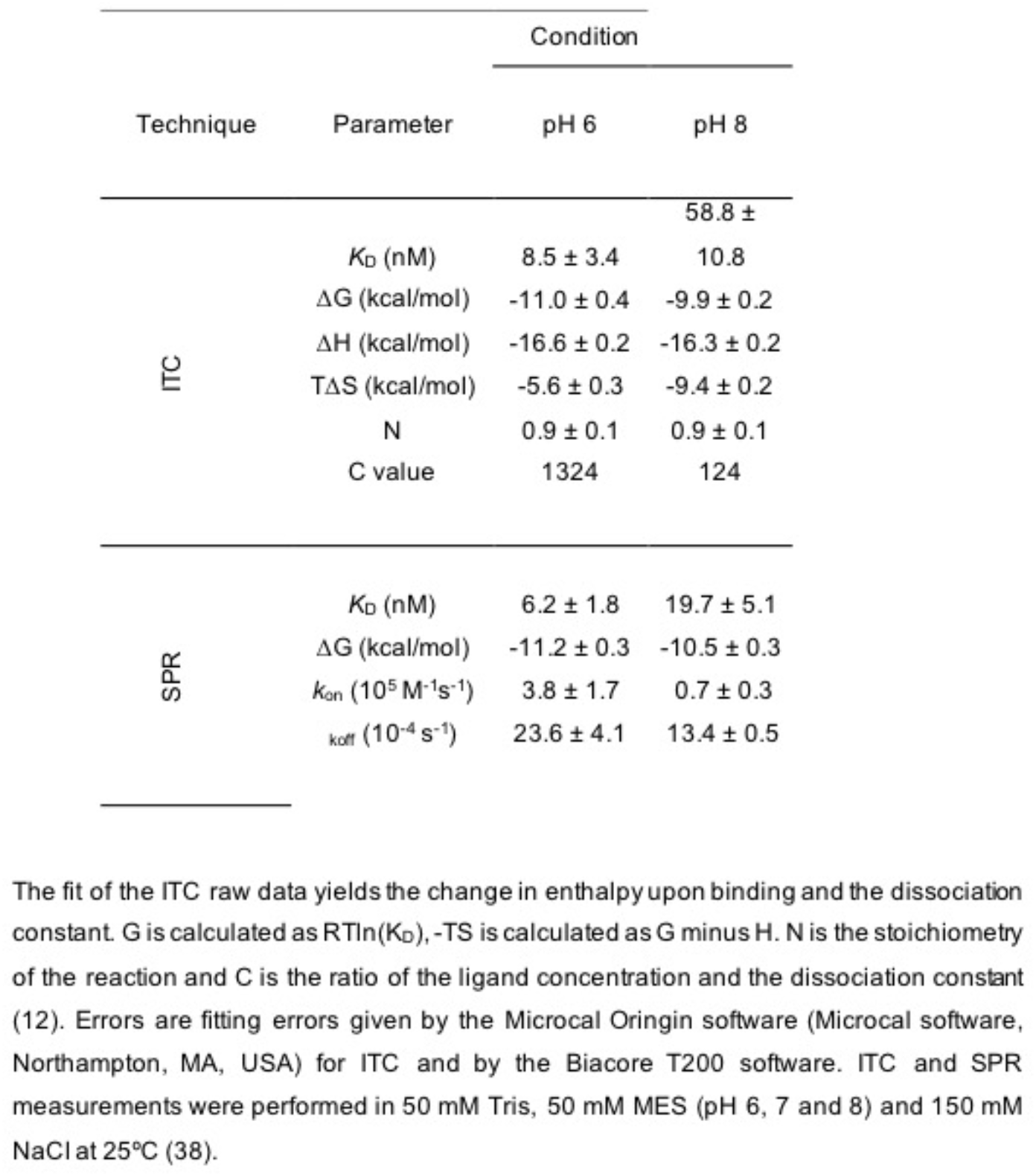
pr-E binding parameters.

### pH-dependent binding of pr and sE

We used size-exclusion chromatography (SEC) combined with multi-angle light scattering (MALS) to analyze the binding of pr to YFV sE at neutral and acid pH. At pH 8.0 and pH 5.5 both pr and YFV sE elute as monomer (Fig. 3A top and bottom panels). Although the sE monomer has a higher molecular mass than the pr monomer, it elutes from the SEC column at a later peak, corresponding to the elution of small molecules. This behavior has been described for other class II proteins (14) and it is probably due to the interaction of the exposed fusion loop with the resin of the column that delays elution. The pr/sE complex at both pH elutes as a 65-60KDa peak (Fig. 3B top and bottom panels) containing both sE and pr proteins as shown by the SDS-PAGE analysis of the peak fractions (Fig. 3C top and bottom panels). This is different from what has been observed for TBE sE protein (7) and for DENV or ZIKV sE proteins (Suppl. Fig. S2). The ZIKV sE protein is a dimer at pH 8.0 (Suppl. Fig. S2Aa top panel) and associates to pr only at pH 5.5 (Suppl. Fig. S2Ab top and bottom panels and Suppl. Fig. S2Ac top and bottom panels). The DENV sE protein at pH 8.0 is a monomer (Suppl. Fig. S2Ba top panel). This monomer dissociates into two peaks at pH 5.5, that we called P1 and P2 (Suppl. Fig. S2Ba bottom panel). We further analyzed only P2 for the complex with pr because P1 was shown to be probably misfolded sE since it was not recognized by EDE neutralizing antibodies (9). Peak P2 showed a retarded elution profile (18ml elution volume, Suppl. Fig. S2Ba bottom panel), similarly to YFV sE protein. After mixing with pr at pH5.5, P2 peak shifts to 14.4ml elution volume (Suppl. Fig.S2Bb bottom panel) suggesting interaction with pr and prevention of the FL interaction with the resin of the column. At pH8.0 instead, the sE and pr peaks overlapped and it was not possible to distinguish whether pr interacts with sE or not (Suppl. Fig. S2Bc top panel).

**Figure 3:**
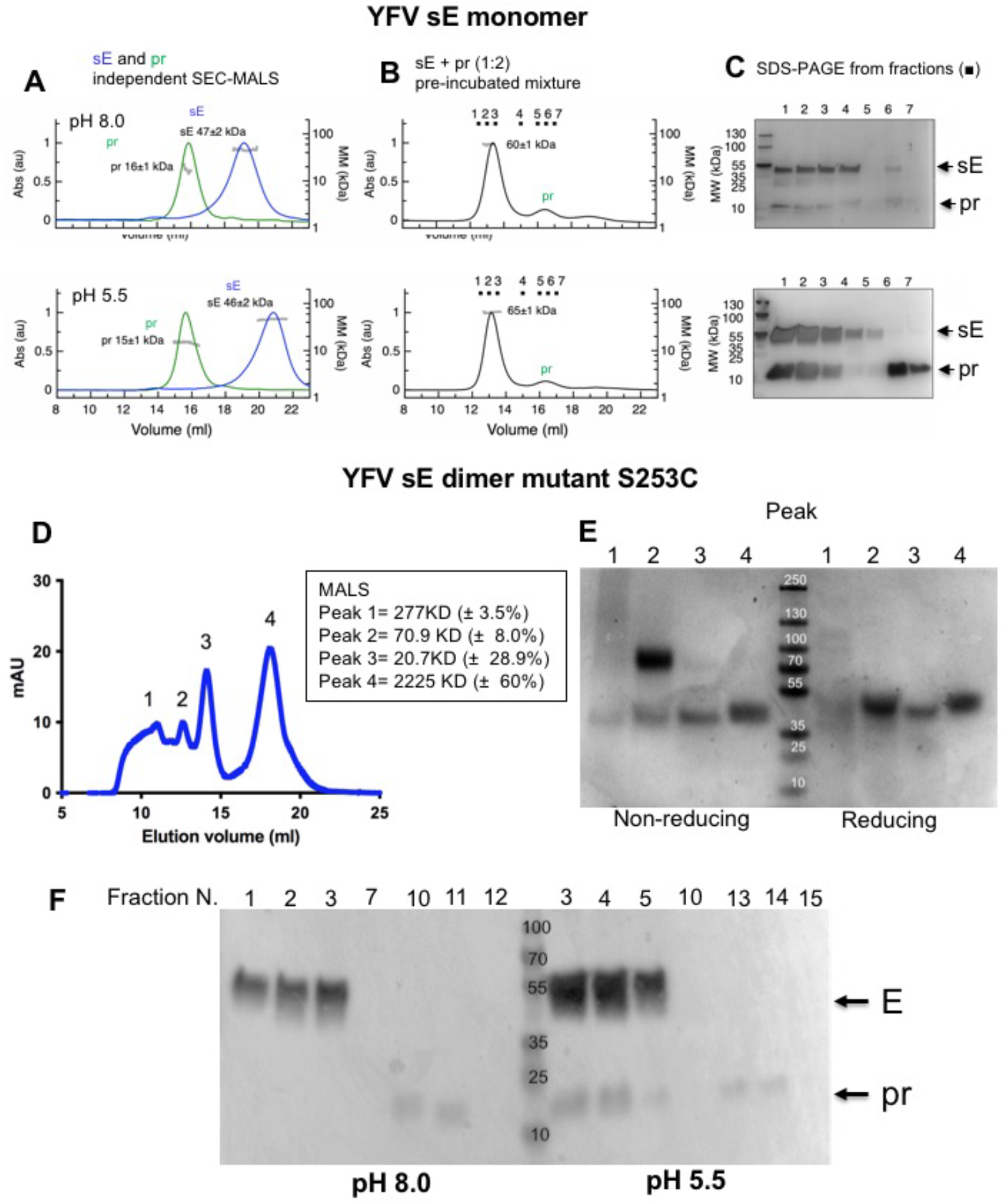
YFV pr protein binds to the E protein monomer at both neutral and acid pH while binding to the E protein dimer is impaired at neutral pH. **(A-B)** SEC-MALS elution volume profiles. Left y axis: the ultraviolet absorbance normalized by setting the highest peak to 1. Right y axis: molecular mass (kDa) determined by MALS, with the values for each species indicated on the corresponding peak. **(A)** Equilibrated SEC-MALS elution profiles of isolated sE (in blue curves) and isolated pr (in green curves) equilibrated at pH 8.0 (top panel) and pH 5.5 (bottom panel). **(B)** SEC-MALS elution profiles of a mixture of sE with pr in excess (1:2 sE:pr monomermonomer molar ratio) at pH 8.0 (top panel) and pH 5.5 (bottom panel). The fractions analyzed by SDS-PAGE in (C) are indicated (1-7). (C) SDS-PAGE and silver nitrate staining of the SEC fractions indicated in (B) at the corresponding pH. (D) SEC elution volume profile of a YFV single cysteine mutant E dimer (S253C) at pH 8.0. The four peaks have been run independently to determine the molecular mass (kDa) by MALS. The MALS results for each peak are listed in the inset. (E) SDS-PAGE Coomassie staining of the four peaks indicated in (D) under reducing and non-reducing conditions. Peak 2 contains the stabilized sE dimer confirming the molecular mass calculated by MALS (70.9 kDa in (D)). **(F)** Western blot of an SDS-PAGE in reducing conditions probed with an anti-Strep antibody of SEC fractions from sE dimer in complex with pr at pH 8.0 and pH 5.5 as indicated and as described in Methods.

Since the E protein is a dimer at the surface of the virus but the sE of YFV is a monomer, we sought to test the interaction of pr protein with a YFV E dimer. To obtain this protein in solution, we engineered a mutation to cysteine in position S253 to induce the formation of a disulfide bond and link the two sE protomers, following the same strategy previously used to stabilize the dengue E dimer (15). The S253C mutant SEC profile showed the presence of high-molecular weight aggregates and peaks corresponding to monomeric protein but a fraction of the protein was produced as a disulfide linked dimer as shown by MALS and SDS-PAGE analysis (Fig. 3D and 3E). Interaction of this dimer with pr resulted in an association only at pH 5.5 (Fig. 3F) similarly to Zika sE dimer and to a stabilized dimer construct for DENV2, mutant A259C (Suppl. Fig. S2Ca,b,c). In conclusion, from the analysis of several mosquito-borne flaviviruses, the pr binding site on the E protein is accessible at low pH on both E monomer or dimer but it becomes hidden on the dimer at neutral pH. However, in the context of the E monomer, the YFV E protein is the only one showing an interaction with pr also at neutral pH (Table 3).

**Table 3.** Summary of sE oligomerization states and pr binding for YFV sE, YFV sE, S253C (noted as YFVd), ZIKV, DENV2 sE A259C (noted as DENV2d) and TBEV sE, at pH 5.5 and pH 8.0. Values for tick-borne encephalitis virus (TBEV) are extracted from (7).

| sE variant | pH 5.5 | pH 5.5 + pr |  | pH 8.0 | pH 8.0 + pr |  |
| --- | --- | --- | --- | --- | --- | --- |
|  | sE Oligomer /MM (kDa) | sE Oligomer /MM (kDa) | sE-pr Binding | sE Oligomer | sE Oligomer /MM (kDa) | sE-pr Binding |
| YFV | Mon. / 46±2 | Mon. / 65±1 | Yes | Mon. / 47±2 | Mon. / 60±1 | Yes |
| YFVd | nd | nd | Yes* | Dim. / 71±8 | nd | No* |
| ZIKV | Aggregation | Mon. / 54±3 | Yes | Dim. / 91±2 | Dim. / 91±1 | No |
| DENV2 | Mon. / 45±1 | Mon. / 48±1 | Yes | Mon. / 51±4 | Mon. / 49±1 | nd |
| DENV2d | Dim. / 92±3 | Dim. / 135±7 | Yes | Dim. / 91±4 | Dim. / 91±2 | No |
| TBEV | Mon. / 52±1 | Dim. / 93±1 | Yes | Dim. / 95±1 | Dim. / nd | No* |

nd, not determined
YFVd is for YFV sE S253C mutant; DENV2d is for DENV2 sE A259C mutant
\*binding determined only by SEC and SDS-PAGE
(see Fig.3 for YFVd and (7) for TBEV)

### YFV pr protein blocks insertion of sE protein into membranes

We tested the effect of the presence of pr on the interactions of sE protein with membranes by measuring co-flotation with liposomes in density gradients (Fig 4A and B). In this assay we mixed purified sE protein with liposomes (see Methods for composition) and, after incubation at neutral or low pH, we separated the complex on a density gradient. If the protein inserts in the membrane of the liposomes, it will co-float to the top fraction of the gradient. We found that at pH 8.0 sE remained at the bottom of the gradient and do not interact with the liposomes in spite of FL exposure (Fig. 4A, left column). Instead, at pH 6.0, about 45% of the sE protein floated to the top fractions (Fig. 4A, B). In the presence of pr, we found a dose-dependent inhibition of sE co-flotation, such that at a molar ratio of 1:1 pr:sE there was no sE protein found in the top fraction, in line with the K_D_ of 10 nM or less of the pr/sE complex at pH 6.0 (Table 2). These results are different to those obtained in the DENV2 system, where a 10-fold molar excess of pr was required to inhibit liposome insertion (13), again indicating that the interaction of pr with the E protein is much stronger in the case of YFV.

**Figure 4:**
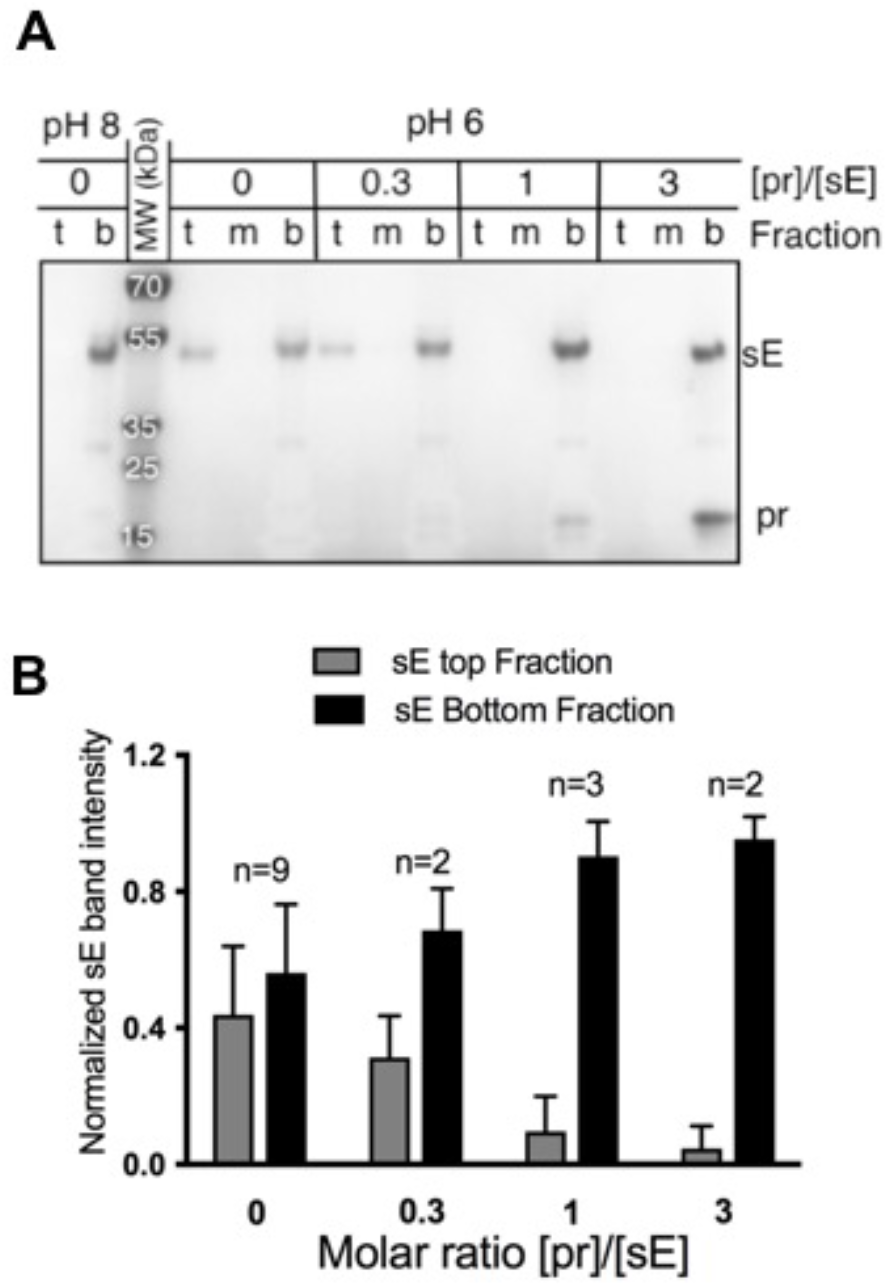
Binding of pr prevents sE insertion into membranes. **(A) Co-floatation assay**. Five µg of purified sE protein was mixed with different amounts of purified pr protein and with liposomes (refer to Methods for lipid composition). After addition of buffer at the indicated pH and over-night incubation at 30°C, the protein-liposomes mixture was separated on an Optiprep gradient. Coomassie stained SDS-PAGE of top (t), medium (m) and bottom (b) fractions is shown. sE protein liposome co-flotation was performed at pH 8.0 (left columns) and at pH 6.0 in presence of pr at sE:pr molar ratios 0.3, 1 and 3 (right columns). **(B) Histogram** of normalized sE band intensity from top and bottom fractions to the amount of sE present in the bottom fraction at pH 8.0. Several flotation assays were included in the calculation using Image J software. Errors are standard deviation calculated from at least two experiments.

### Interaction of pr with the YFV viral particle inhibits viral fusion

Viral fusion to membranes can be measured using lipid mixing fusion assays. We used a system based on fluorescence resonance energy transfer between the fluorophores 7-nitro-2-1,3-benzoxadiazol-4-yl (NBD) and rhodamine covalently coupled to lipids. The fluorescence is quenched by a high concentration of the two fluorophores in the liposomes, and becomes de-quenched upon dilution into the lipids derived from the viral membrane upon fusion of the two lipid bilayers, allowing to follow the lipid merger reaction. The fluorescence profile observed upon mixing YFV strain 17D virus with the NBD/rhodamine labeled lipids at different pH values is displayed in Fig. 5A. Fluorescence dequenching is optimal between pH 5.6 and pH 6.2 and is negligible at neutral pH. A plot of the mean intensities reached at each pH shows a peak of lipid mixing at around pH 6.0 (Fig. 5B). We therefore used pH 6.0 to test the inhibition of lipid-mixing by recombinant pr added at different pr:E stoichiometries to the virus preparation before mixing with liposomes and found a dose-dependent inhibition of the reaction by exogenous pr (Fig. 5C). For the fusion experiments, we used YFV17D virus because of safety reasons, since the vaccine strain can be manipulated under BSL2 conditions. The vaccine strain 17D carries 10 amino acids mutation in the sE protein (16) but their localization does not interfere with the pr/sE binding site. We quantified the relative stoichiometry of pr:E by western blot as described in the Materials and Methods section (see Suppl. Fig. S3). Differently to the results observed on the inhibition of YFV sE protein insertion into liposomes by pr (Fig. 4A), we observed a requirement of pr in excess of at least 10 times to obtain 100% inhibition of lipid mixing (Fig. 5D). This discrepancy suggests a different affinity of pr for E on virions compared to sE in solution. This is probably due to the different accessibility of the pr binding site in the context of the E dimer (present on the virus) compared to the E monomer present in solution. The pr binding site could be indeed buried in the E dimer of the viral particle at neutral pH and become available only when the dimer is opening at low pH. To test this hypothesis, we mixed pr with YFV particles in an excess of 50:1 pr:E stoichiometry at various pH values, and measured the amount of pr brought down upon pelleting of the virion by ultracentrifugation (Fig. 5E). This experiment showed very little pr co-precipitating with the virus at pH 8.0, and a maximum of co-precipitation at pH6.0, suggesting a pH dependent exposure of the pr binding site on virions (Fig. 5F).

**Figure 5:**
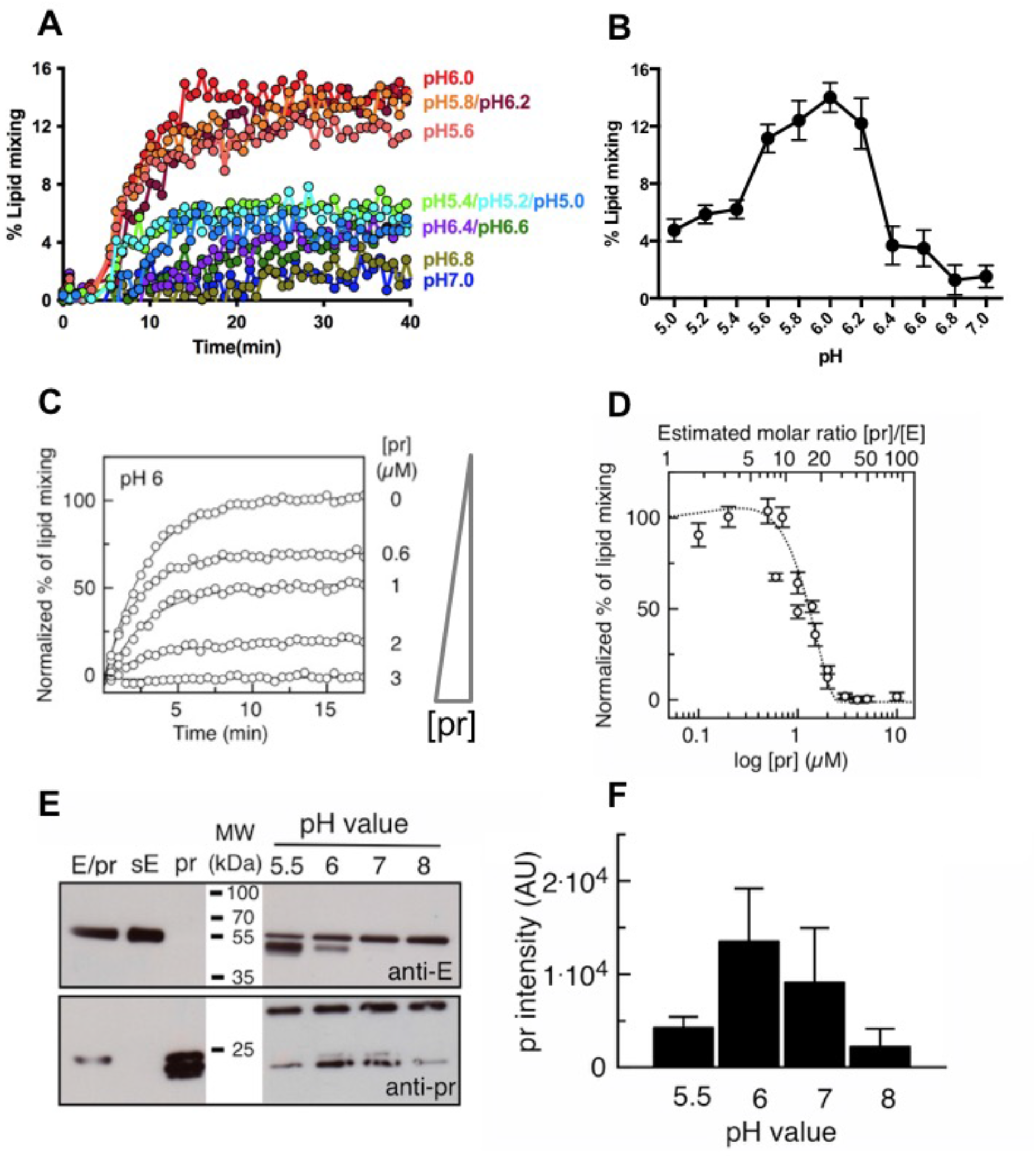
Effect of pr binding to the E protein of the viral particle. **(A) Lipid mixing assays** between YF17D virus and NDB/Rho-labeled liposomes recorded from pH 5.0 to pH 7.0 at every 0.2 pH units. About 107-108 ffu of purified virus was mixed with 500nM labeled liposomes resuspended in buffer at different pH. Fluorescence emission was recorded for 40min in a multiplate reader fluorimeter (Tecan M1000) and the reaction was stopped by addition of detergent to measure the maximal (100%) signal. The represented extent of lipid mixing was related for each pH to the maximum signal recorded upon lipid dilution by detergent addition (see Methods). **(B) Plot of mean fluorescence signal** registered at min. 7-37 for each pH. **(C) Representative curves of normalized NBD fluorescence intensity** recorded at 535 nm as a function of pr concentration, lines are mono-exponential fits to the data. Sample without pr was considered 100%. About 107-108 ffu of purified virus was mixed with increasing amount of purified pr and incubated at 37°C for 30min. Virus/pr mixture was then added to 200nM liposomes in MES buffer pH 5.5. The final pH of the mixture was pH 6.0. Fluorescence emission was recorded for 30min in a plate reader fluorimeter (Tecan M1000) and the reaction was stopped by addition of detergent to measure the maximal signal. **(D) Percentage of lipid mixing** as a function of pr concentration. Top x-axis corresponds to the [pr]/[E] molar ratio as estimated by western blot (see Methods and Suppl. Fig.S4). The dashed line is a guide to eye. **(E) Binding of exogenous pr to YFV viral particle**. Western blot with E-or pr-specific antibodies. About 108 ffu of virus was mixed with an excess (1:50) of exogenous purified pr protein and incubated in buffer at different pHs. The complex was then pelleted by ultracentrifugation and analyzed by SDS-PAGE and western blot. **(F) Histogram** representing the values of pr band intensity from two experiments.

### The YFV sE dimer

While the YFV sE protein is mainly a monomer in solution, we were able to obtain crystals of a sE dimer using the construct without prM. This protein formed tetragonal crystals that diffracted to 3.5Å (Suppl. Table S1). The structure, determined by molecular replacement (using the 6EPK structure) and refined to a free R factor of ∼ 27% (see Methods), showed the typical head-to-tail sE dimer conformation observed initially for sE of TBEV (17) and later for the DENV2 (18), JEV (19), and ZIKV (20), (21) counterparts. There are two main sE dimer interfaces, the first by the dimer axis and the second one involving the fusion loop, away from the dimer axis. The first interface involves antiparallel interaction of the polypeptide chain around helix *α*B (Fig. 6A), including several inter-protomer hydrogen bonds, some of which involving main-chain / main-chain interactions. In the second interface, the FL at the tip of domain II packs against domains III and I of the other protomer in the dimer (Fig. 6B). The FL residue Trp101 has its side chain covered by that of Lys308 of domain III, while the FL main chain is partially tucked in between two short helices in domain I, the N-terminal helical turn (N-helix in Fig. 6B) and the “150-helix” (150-loop forming an *α* helix) described in more detail below (Fig. 6B).

**Figure 6:**
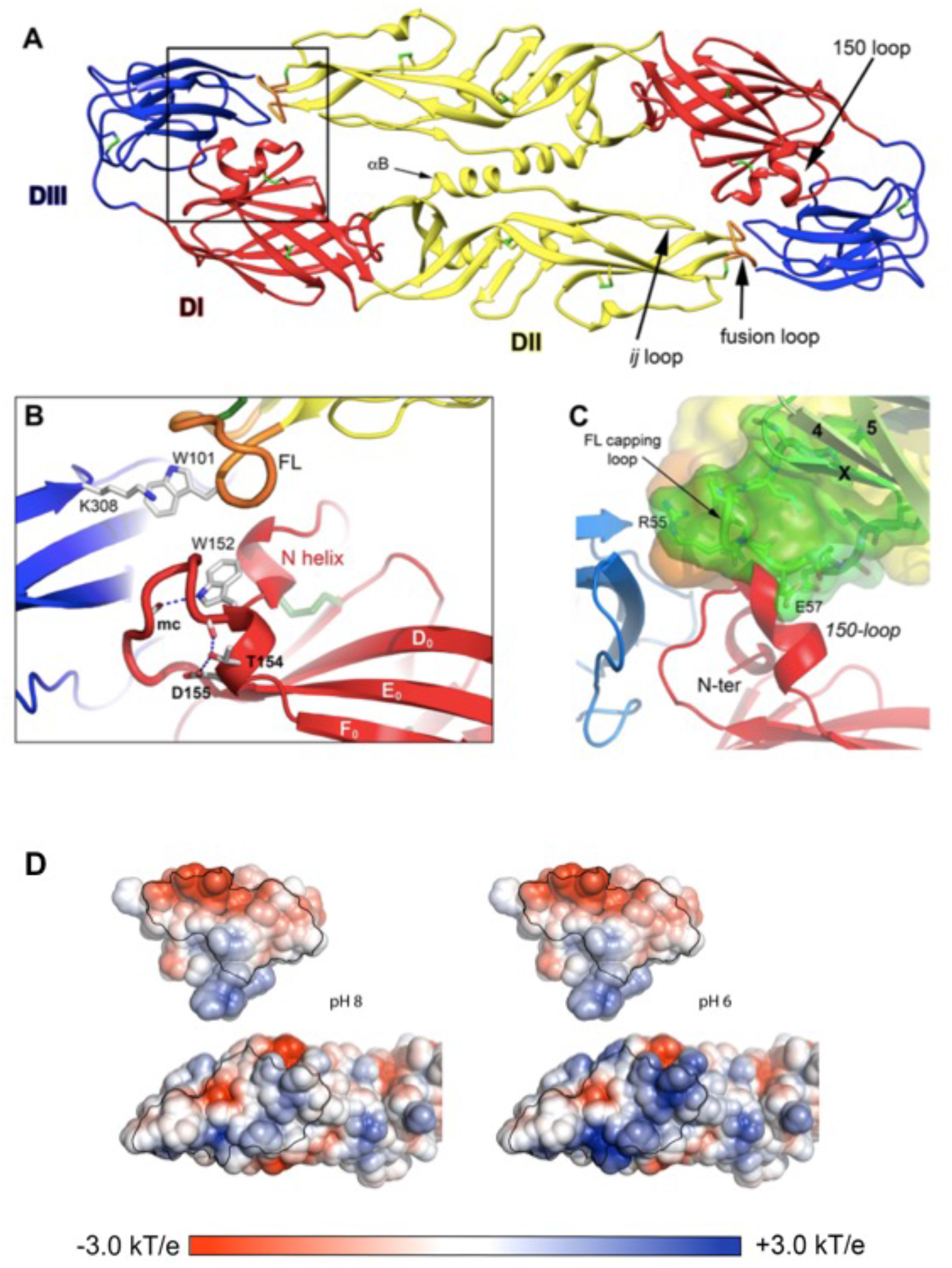
YFV sE dimer and structural analysis of its interaction with pr protein. **(A) Ribbon view** of the crystallographic YFV sE dimer colored according to definition described in figure 1. The fusion loop in orange is buried at the dimer interface against 150-loop and *ij*-loop, as indicated. **(B) Close-view** showing the proximity of the 150-loop with the N-terminal of E. **(C) Close-view** of modeled binding of pr protein on the sE dimer showing the clash of the pr capping loop with the 150-loop and N-terminal of E. The superposition of pr/sE structure on sE dimer was done using the tips of E-domain II containing the fusion loop (β-strands *b,c,d* and *ij;* see Figure 1) of pr/sE monomer (6EPK). **(D) Electrostatic potential surfaces of pr/sE complex**. The electrostatic potential surfaces are displayed as an open book representation of the pr/sE interaction computed at two pH: pH 8.0 (left) and pH 6.0 (right). The potential of pr (top views) is not dependent of the pH while the potential of E (bottom views) appears to direct the change of the electrostatic at the interaction surface with pr.

### The 150-loop

The YFV sE dimer displays a unique organization of the 150-loop in domain I, which is highly variable in sequence across flaviviruses and connects *β*-strands E_0_ and F_0_ in domain I (Fig. 6B). Most flaviviruses carry an N-linked glycan at positions 153 or 154, except for YFV, for which only a few attenuated strains are N-glycosylated (22). The “150-helix” (residues 149-155) is highly exposed at the dimer surface. A short helix in the 150-loop is present in other flaviviruses as well (MBEVs and ZIKV) albeit oriented almost at 90 degrees (7). The side chain of Trp152 appears as an important element of the “150-helix”, as it packs against the N-terminal end of the polypeptide chain, which is buried underneath. The positively charged N-terminal Ala1 is neutralized by a salt bridge and hydrogen bond with the Asp42 sidechain, which is also buried. The buried N-terminal end of the protein appears to confer a specific structure to domain I, as in the structure of the pr/sE complex of dengue virus serotype 2, in which a linker connected the region of prM just upstream of the trans-membrane (TM) segment to the N-terminus of sE (thereby by-passing the TM region), showed a disordered 150-loop with the N-terminal helix continuing in the linker and projecting out at the top of domain I (7). The N-terminus of the wild type YFV E protein indeed participates in a network of hydrogen bonds also involving residues from domain III.

The first helical turn of the “150-helix” is somewhat distorted, but the second turn is further constrained by a hydrogen bond between the side chains of the consecutive Thr154 and Asp155 (Fig. 6B). Importantly, one of the virulence determinants of YFV in a hamster model was found to map to position 154, which was identified as conferring virulence when Thr154 was replaced by Ala (23). In that study, mutating Asp155 to Ala resulted in a variant with the same virulence phenotype even when Thr154 was maintained, suggesting that the hydrogen bond between these two adjacent side chains residues is important for stabilizing the relevant conformation required for interactions with the host that affect virulence. In summary, the “150-helix” is highly exposed and structured at the dimer surface, in a region important for stabilizing interactions between domains I and III and the fusion loop on the adjacent dimer subunit.

To understand the interactions of pr with the sE dimer, we modeled the pr binding site (as determined in the pr/sE monomer (6EPK)) on one subunit of the sE dimer and identified a clash between the pr capping loop and the E 150-loop. This clash suggests an impaired binding unless the 150-loop moves out of the way in an “open” position as it has been shown for TBE dimer at low pH (7). To understand which interactions would allow the pH-dependent movement of the 150-loop and release of pr at neutral pH, we analyze the electrostatic potential of E and pr surfaces at their binding site at pH 8.0 and pH 6.0 (Fig. 6D). We could still detect a fair charge complementarity at pH 8.0 which can explain why the affinity of pr for the E monomer is still high at neutral pH. It remains to be determined if, in the context of the dimer, these interactions are sufficient to expel pr at neutral pH or if additional re-arrangements of the envelope proteins are required.

## DISCUSSION

Our data provide a structural and functional analysis of the interaction between pr and E protein of yellow fever virus. Comparison of these data to other flaviviruses, such dengue and Zika viruses, show a general mechanism of action of pr in protecting the FL at low pH, a critical step of virus maturation. We show that pr associates to the E dimer at low pH for YFV, DENV and ZIKV but this interaction is lost at neutral pH. However, only the YFV E monomeric protein showed an interaction with pr also at neutral pH. This interaction is stabilized by several inter chain contacts that are absent in the other flaviviruses (see Table 1). We show, as previously reported in the literature, that exogenous addition of purified pr to sE interferes with its insertion into liposomes at low pH in a floatation assay, an assay mimicking the dimer-to-trimer transition occurring during viral fusion (4), (13). Differently from what it has been shown for DENV, where high concentration of pr were required to inhibit sE co-floatation with the liposomes, a 1:1 pr:sE molar ratio was sufficient for YFV sE to block membrane insertion, confirming the high affinity of these two proteins. Moreover, we were able to show, using infectious virus in a fusion assay, that this interaction actively blocks viral fusion. In contrast to the results obtained with the purified protein, we needed a 10-fold excess of pr protein to completely inhibit fusion of infectious virus. This is due to the fact that the E protein at the surface of the virus is present as a dimer and, at neutral pH, the FL is not accessible to pr as it is on the monomeric purified protein. In both experiments, the pr/E complex was generated at neutral pH and after addition of the liposomes the pH was lowered to the chosen acidic value. Virus and purified sE protein at low pH, in absence of membranes, would indeed aggregate interfering with the read-out of the assay. These results confirm what we have observed in our SEC-MALS analysis that showed pr/sE binding at neutral pH only for the monomeric form of sE and not for the dimer. Our MALS analysis revealed also some difference in the way flaviviruses handle the pr/E interaction. The FL is protected by the pr interaction at low pH but while ZIKV dimer dissociates at acidic pH, TBE remains dimeric. DENV and YFV instead are monomeric at both neutral and acidic pHs (Table 3). These data support how different flavivirus sE proteins vary regarding their pH sensitivity to dimerization/dissociation, while the molecular mechanism dictating pr binding/unbinding and thus flavivirus maturation, is common to all flaviviruses.

During maturation there are three critical steps in which is mandatory for flaviviruses to protect the fusion loop from premature membrane insertion. First, after budding in the ER, the immature virus carries pr bound to the FL on top of the trimeric (prM/E)_3_ spikes; second, during the transit through the acidic TGN, the pH-induced trimer-to-dimer transition generates immature smooth particles carrying pr on top of the FL exposing the furin cleavage site; third, after pr-M cleavage, at the neutral pH of the extracellular milieu, pr is displaced from the FL by the snap-lock movement of the 150loop (7). While pr binding to FL is the key interaction throughout these steps, its regulation occurs via combined action of several regions of the pr/E complex, identifying the pr-binding site as a leading character in the flavivirus maturation process. Not surprisingly this region is targeted by highly cross-neutralizing antibodies (20). Our structure of the YFV sE dimer confirmed the folding previously described (24) and showed how the 150 loop at neutral pH is in closed conformation and would clash with pr binding, a mechanism previously described for TBE (7). This explains the higher ratio of pr required to block viral fusion in our lipid mixing experiments.

In conclusion, we describe the molecular interactions regulating a crucial process in flavivirus maturation. Interestingly, while the basic organization of the interactions is common to all flaviviruses, each virus seems to modulate them differently. In particularly, we found for yellow fever a stable association with pr also at neutral pH suggesting that its release from the mature particle cannot occur exclusively by a passive pH-dependent change of charges but it will require an active reorganization involving the viral particle in its whole.

## METHODS

### Recombinant pr/sE protein production

The YFV Asibi pr/sE and sE constructs were cloned onto a pMT-derived vector (25). This vector allows expression of the gene of interest downstream an insect signal peptide BiP and in frame with an enterokinase or a thrombin cleavage site followed by a StrepTag, for purification purpose. The sequence encoding for prM and the ectodomain of E for Asibi strain (NCBI AY640589) was taken from pACNR-113.16 (Rice and Barba-Spaeth, unpublished). Single cysteine mutation S253C was introduced to generate a disulfide stabilized E dimer protein. All the constructs were restricted to residues 1 to 392 for E. *D. melanogaster* S2 pseudo-clonal pools were generated by co-transfection with a pCoPURO (26) by Effectene transfection (QIAGEN). For expression, cells were induced at a density of 1×10^6^ cells per mL with 500 µM Cu_2_SO_4_ for 10 days or 5 µM CdCl_2_ for 7 days. The supernatant was then harvested, concentrated on a Vivaflow 200 concentration system with a 10 kDa-cutoff membrane (Sartorius). The pH of the concentrated supernatant was adjusted to 8.0 with 100 mM Tris HCl and avidin was added to a final concentration of 1 µg/mL. Soluble YFV sE protein was then captured on a StrepTactin column, washed and eluted with binding buffer (100 mM Tris pH 8.0, 150 mM NaCl, 1 mM EDTA) supplemented with 2.5 mM desthiobiothin. The peak obtained by affinity chromatography was further purified by a size-exclusion chromatography, using a Superose6 16/300 column (GE Healthcare) with 20 mM Tris HCl pH 8.0 and 150 mM NaCl. The purified protein was then dialyzed against 10 mM Tris HCl pH 8.0 and loaded on MonoQ 5/15 column (GE Healthcare) to be eluted using a step gradient of 240 mM and 400 mM NaCl in the same buffer.

To denature the complex pr/sE under non-reducing conditions, 8 M urea was added to the solution and the two proteins (47KDa and 10Kda) were separated by a size exclusion chromatography (SEC) in 10 mM Tris-HCl pH 8, 6 M urea and 1 M KSCN. Samples were collected and dialyzed overnight against 20 mM Tris-HCl pH 8 to remove any trace of urea. A final purification on a Superdex 200 16/60 in 20mM Tris-HCl pH 8.0 and 150mM NaCl was done to obtain pure and refolded E protein and pr peptide.

ZIKV sE (strain PF13), DENV2 sE (SG strain) and DENV2 sE A259C mutant (16681 strain) were produced as described earlier. Briefly, sE genes with a tandem C-terminal strep-tag in pMT/BIP/V5 plasmid were expressed in *Drosophila* S2 cells (Invitrogen) as described previously (20), (9). Protein expression was induced by the addition of 5 μM CuSO4 or CdCl_2_. Supernatants were harvested 8–10 days post-induction, and sE were purified using Streptactin columns (GE) according to manufacturer’s instructions. This affinity chromatography step was followed by size exclusion chromatography using Superdex 200 10/300 GL column equilibrated in 50 mM Tris (pH 8) and 500 mM NaCl. Pr proteins from YFV, ZIKV (PF13 strain) and DENV2 (16681 strain) were expressed similar to sE proteins, using same pMT/BIP/V5 plasmid with double C-terminal strep tag in were expressed in *Drosophila* S2. Pr proteins were purified using a streptactin columns based affinity step and followerd by a single SEC step using Superdex 75 10/300 GL column equilibrated in 50 mM Tris (pH 8) and 300 mM NaCl.

### Crystallization

#### pr/sE Asibi crystallization

After optimization, crystals diffracted up to 3Å resolution but an analysis of the intensity distribution revealed that the datasets was perfectly and merohedral twinned with apparent space group P4_1_22. To overcome the problem an additional purification step using denaturation / renaturation of the heterodimer under non-reducing conditions was introduced. The reassembled pr/sE complex was concentrated to 3 mg/mL in 20 mM Tris-HCl pH 8 and 150 mM NaCl and crystallized into 100 mM Tris HCl pH 8 and a range of 1.2-1.8 M Li_2_SO_4_. For cryoprotection, crystals were soaked in the precipitation solution plus 25% glycerol and flash-frozen under liquid nitrogen.

#### sE Asibi dimer crystallization

Asibi sE, produced without co-expression of prM, and purified by SEC in 20 mM Tris pH 8, 150 mM NaCl, was adjusted to a concentration of 3.2 mg/mL, and formed highly regular crystals in 1.26 M (NH4)_2_SO_4_ and 0.1 M HEPES pH 7.5. The crystals diffracted at very low resolution and optimized conditions allowed to grow bigger crystals which gave diffractions ranging from 6 Å to 3.7 Å.

### Data collection, Refinement and Model building

Diffraction data were collected at the beamlines Proxima-1 and Proxima-2 at the SOLEIL synchrotron and ID23-1 at the ESRF synchrotron, were processed using XDS package (27) and scaled with AIMLESS (28). Only the diffraction data of the sE dimer crystal shown significant anisotropy. Therefore, this data was elliptically truncated and corrected using the DEBYE and STARANISO programs (developed by Global Phasing Ltd) using the STARANISO server (29). The unmerged protocol applied to this data produced a best-resolution limit of 3.48Å and a worst-resolution limit of 4.87Å with a surface threshold of 1.2 of the local I/σ(I). This corrected data was used for refinement of the sE dimer structure. The structure of Asibi pr/sE (6EPK) was first determined by molecular replacement with the program AMoRe (30) using the atomic models TBEV sE protein (PDB entry 1SVB, 43.4% sequence identity, (17) and DENV pr protein (PDB entry 3C5X, 34.6 % sequence identity, (11). Then, the Asibi sE protein from the pr/sE structure was used as a template for molecular replacement for solving the sE dimer structure. The two models were subsequently modified manually with COOT (31) and refined with BUSTER-TNT (32), (33) or PHENIX.REFINE (34). Refinement was constrained to respect non-crystallographic symmetry and target restraints (35) using high resolution structures of parts of the complexes, as detailed in the Table SUPP 1. TLS refinement (36) (parameterization describing translation, liberation and screw-motion to model anisotropic displacements) was done depending on the resolution of the crystal. The final models of pr/sE (PDB 6EPK) and sE dimer contain all amino acids of YFV sE (1-392) and residues 1 to 80 of pr. Data collection and refinement statistics as well as the MolProbity (37) validation statistics for all the two structures are presented in the Table SUPP 1. The figures of the structures were prepared using the PyMOL molecular graphics system (Schrodinger)(pymol.sourceforge.net).

### Multi-angle static light scattering-Size exclusion chromatography

MALS studies were performed using a SEC Superdex 200 column (GE Healthcare) previously equilibrated with the corresponding buffer, see below. SEC runs were performed at 25 °C with a flow rate of 0.4 mL/min, protein injection concentration was 100 μg. Online MALS detection was performed with a DAWN-HELEOS II detector (Wyatt Technology, Santa Barbara, CA, USA) using a laser emitting at 690 nm. Online differential refractive index measurement was performed with an Optilab T-rEX detector (Wyatt Technology). Data were analyzed, and weight-averaged molecular masses (Mw) and mass distributions (polydispersity) for each sample were calculated using the ASTRA software (Wyatt Technology). For each virus, equilibration buffers for addressing the effect of pH for sE, pr and the sE:pr complex were the three-component buffers, 100 mM Tris-HCl, 50 mM MES, 50 mM sodium acetate and 150 mM NaCl, at pH 5.5 or pH 8.0. The sE:pr complex, in 1:2 molar ratio (monomer:monomer molar ratio), were prepared by incubation in the corresponding three-component buffers. Buffer exchange was performed by extensive dialysis of the sample, 12 h stirring at 4 °C and two 500 mL buffer replacement in 10 kDa molecular weight cut-off dialysis membranes (Spectrum). SEC fractions of sE:pr complexes at pH 5.5 or 8.0 were further analyzed by Coomassie blue or Silver nitrate SDS-PAGE or by western blot using an anti-strep antibody for simultaneously detection of both E and pr proteins.

### Liposomes preparation

Liposomes used for lipid mixing and co-flotation assays were prepared by following a modified film-hydration protocol (38). Briefly, chloroform solutions of DOPC, DOPE, SM, Cholesterol, NBD-PE and Rho-PE, were pooled using glass graduated syringes (Hamilton) in borosilicate tubes at a molar ratio of 1:1:1:3:0.1:0.1, respectively, and a total lipid concentration of 10 mM. The fluorescent lipids (NBD-PE and Rho-PE) were omitted in the preparation of liposomes for co-floatation assays. The organic solvent was evaporated in the tube under a steam of N2 gas yielding a thin lipid film which was further dried by Speed-Vac (Thermo Electron, RVT400), 1 hour at room temperature. The lipid film was resuspended in 20 mM HEPES pH 7, 50 mM NaCl degassed buffer, by vortexing in presence of 180 μm acid washed glass beads (Sigma). The resulting opaque solution, composed by multilamellar vesicles, was subjected to 10 cycles of liquid N2 flash freeze-thaw and extruded using a polycarbonate filter of 100 nm pore size until translucency, more than 20 extrusion cycles. The hydrodynamic diameter and homogeneity of the sample was controlled by dynamic light scattering. The final lipid concentration was determined by a using NBD-PE absorbance at 460 nM and a standard curve. The liposomes were stored under N2 (gas) for up to three weeks at 4ºC. All the lipids as well as the extrusion system were purchased from AVANTI Polar Lipids (USA). Abbreviations: DOPC: 1,2-dioleoyl-sn-glycero-3-phosphocholine; DOPE: 1,2-dioleoyl-sn-glycero-3-phosphoethanolamine; SM: Sphingomyelin (brain, porcine); NBD-PE: 2-dioleoyl-sn-glycero-3-phosphoethanolamine-N-(7-nitro-2-1,3-benzoxadiazol-4-yl); Rho-PE: 1,2-dioleoyl-sn-glycero-3-phosphoethanolamine-N-(lissamine rhodamine B sulfonyl) (ammonium salt).

### sE-liposomes co-floatation assay

Renatured sE and pr proteins were mixed at different molar ratio and incubated for 10 min at RT before addition of liposomes. The mixture was further incubated for 10 min at RT before overnight incubation at 30°C under acidic conditions. The liposomes were then separated by ultracentrifugation on an Optiprep (Proteogenix 1114542) continuous 0-30% gradient. Aliquots from top and bottom fractions were analyzed by Coomassie gel or by western blot gels using in house produced anti-YFV E (E21.3) mouse monoclonal antibody. At least two and up to nine experiments were performed for the different molar ratios tested, the bands intensity from top and bottom fractions were analyzed by ImageJ software and plotted as ratio to total protein present in each floatation assay.

### Isothermal titration calorimetry (ITC)

We titrated 10 µM of E in the cell with several injections of 100 µM pr. The injection volume was 2 µL. We continued the injections beyond saturation to determine the heat of ligand dilution, which was subtracted from the data prior to fitting with a single site binding model. We used Microcal ITC200 from Microcal and the associated Origin software for fitting of the data. The two-component buffer was prepared by dissolving appropriate weights of each component in water (39). The resulting solution had a pH of 8.3, which was taken to the desired value with concentrated HCl. The pH was measured in a Sartorius PB11 pH-meter. The protein samples were extensively dialyzed prior the titrations. ITC measurements were performed in 50 mM Tris, 50 mM MES (pH 6, 7 and 8) and 150 mM NaCl at 25ºC (Table 2).

### Surface plasmon resonance (SPR)

The affinity of the sE protein for the pr peptide was measured by SPR using a Biacore T200 system (GE Healthcare Life Sciences) equilibrated at 25ºC. The carboxylic groups of a Series S CM5 sensor chip were activated for 10 min using a mix of N-Hydroxysuccinimide (NHS, 50 mM) and 1-ethyl-3-[3-(dimethylamino)propy1]-carbodiimide (EDC, 200 mM). The Strep-Tactin XT (IBA lifesciences) at 2 µg/mL in acetate pH 5 was injected for 20 min, followed by deactivation with 1 M ethanolamine for 7 min, reaching a density of 800 resonance units (1 RU corresponds to about 1 pg/mm2) of amine coupled Strep-Tactin XT. At the start of each cycle, double strep tagged sE protein was captured on a Strep-Tactin XT surface for 3 min at 5 µg/mL. Eight concentrations of pr peptide (2-fold dilutions ranging from 100 nM to 0.78 nM) were then injected at 30 µl/min for 600s. At the end of each cycle, the surfaces were regenerated by sequential 15s injections of Gly-HCl pH 1.5 and 10 mM NaOH. Experiments were performed in duplicate, using 3 different running buffers, 50 mM MES, 50 mM Tris (pH 6, 7 and 8) with 150 mM NaCl and 0.2 mg/mL BSA at 25°C (Table 2). The association and dissociation profiles were fitted globally using the Biacore T200 evaluation software (GE Healthcare) assuming a 1:1 interaction between sE and pr.

### Virus stocks

Yellow fever 17D (YF17D) viral stocks were derived from pACNR/FLYF plasmid (40) containing the full length infectious YF17D-204 genome under a SP6 promoter, after electroporation of in vitro-generated RNA transcripts in SW13 cells as previously described (41). Briefly, 3 μg of RNA were mixed with 4 × 10^6^ SW-13 cells in PBS and pulsed in 2-mm-gap electroporation cuvettes (BTX) with an electroporator (BTX Electro Square Porator model T820) set for 3 pulses at 800 V with a pulse length of 60 μs. After a 10-min recovery phase at room temperature, cells were plated in a p75 flask in complete medium (Minimum Essential Medium supplemented with 10% fetal bovine serum (FBS), 1 mM sodium pyruvate, 2 mM Glutamax and 0.1 mM MEM non-essential amino acids). Virus stocks were harvested 48h post-transfection with typical yields of 10^7^-10^8^ FFU/mL as determined by focus forming assay on SW13. Single use aliquots were stored frozen at -80°C until use.

### Virus purification

SW13 cell monolayers were infected at low MOI (0.1 ffu/cell) and supernatants were collected 48h post-infection. YF17D virus was recovered by precipitation with 8%(w/v) PEG 8000 for 1h at 4ºC and purified on a step tartrate-glycerol gradient (40-10%(w/v) tartrate - 5-30%(w/v) glycerol) by over-night ultracentrifugation in SW41 at 30 Krpm. Virus band was recovered by needle puncture at the side of the tube and virus titers were determined by focus forming assay. The total amount of virus present in the preparation was quantified by comparison against known amount of purified sE protein and western blot with YF E-specific antibody E21.3. The virus band buffer corresponded to about 25%(w/v) tartrate, 15%(w/v) glycerol and 0.02%(w/v) BSA. The virus preparation was kept at 4°C until use. Buffer-alone gradients were run and collected in parallel to each virus preparation to be used as blank in the functional assays.

### Focus-forming assay

Serial dilutions of the virus preparations (1/10) were prepared in 1% FBS / PBS. Each dilution was added to SW13 cells and foci were developed in the presence of 1,5% methylcellulose for 2 days in 96 well plates. Foci development was stopped by fixation with 4% formaldehyde and foci were then stained using a mouse-anti-NS1 antibody (1A5) (gift from Jacob Schlesinger, Rochester University) and a horseradish peroxidase (HRP) conjugated secondary anti-mouse antibody (ThermoFisher 31430). The foci were visualized by diaminobenzidine (DAB) (Sigma D5905) staining and imaged using the ImmunoSpot S6 Analyser (Cellular Technology Limited).

### pH triggered lipid mixing, pH and pr titrations

We adapted, from standard lipid mixing assays protocols using the pair of probes NBD-PE and Rh-PE (42), (43), a pH-triggered assay to monitor the effect of pH or pr on the extent of lipid mixing between YF 17D virus and labelled liposomes. Mixture reaction for pH titrations: 10 µl of purified virus (10^9^-10^10^ ffu/mL) were added to 100 µl of 500 nM labelled liposomes diluted into 300 mM citrate-phosphate buffer at pH 5.0, 5.2, 5.4, 5.6, 5.8, 6.0, 6.2, 6.4, 6.6, 6.8 or 7.0. Mixture reaction for pr titrations: 10 µL of purified virus (10^9^-10^10^ ffu/mL) were incubated in a multi-well plate (Greiner) with increasing amounts of purified pr protein for 30 min at 37ºC in 100 mM Tris HCl pH 7.5 and 150 mM NaCl, 50 µL total volume. Subsequently, 100 µl of 200 nM NBD-PE and Rho-PE labelled liposomes in 50 mM MES pH 5.5, was added to the virus/pr complex using a multichannel pipette and gently mixed three times prior data collection (average dead time 40s). The pH after the mixture was 6.0±0.2. For both titration assays, the emission fluorescence of NBD was recorded in a multi-plate reader fluorimeter (Tecan M1000), with an excitation and emission wavelength of 460 nm and 539 nm and slits widths of 10 nm and 20 nm, respectively, during more than 3 times the end of the lipid mixing reaction (∼10 minutes) at 25ºC. The maximum NBD emission signal was recorded by addition of 10 µl of 2.5% C13E8 (Polyoxyethylene(8)tridecyl Ether, Anatrace) for 10 minutes. For pr titrations, a mock reaction (no virus) was performed for each pr concentration by using the same virus buffer and used as reference signal. The extent of lipid mixing was calculated from the recorded intensities (I) by (I-I0)/(I100-I0), with I0 the initial intensity and (I100) the maximum NBD emission signal recorded upon addition of detergent. The % of lipid mixing was calculated by the end point parameter of the fitting of the data to a mono exponential equation using ProFit software (QuantumSoft). For pr titrations, we normalized all the curves to the % of lipid mixing measured in absence of pr. The concentration of viral E protein in the final volume of the assay was quantified by western blot. Shortly, a range of 25 to 200 ng of recombinant E was used as a standard curve and in the same SDS PAGE gel we loaded 0.15 µl to 10 µL of purified virus. The western blot was revealed with the E21.3 antibody. Bands intensities were calculated in ImageJ software (44) and used to interpolate the amount of E in the virus to a standard curve of purified E protein (25-200ng) by linear regression. The concentration of viral E of 50-80 nM was used to refer the titrated concentrations of pr as a pr/E molar ratio.

### Co-precipitation of purified pr with YF 17D virus

Cell culture supernatant containing 10^8^ total particles of YF 17D virus was pelleted over a 20% sucrose cushion and resuspended in 100ul of TNE buffer (10 mM Tris pH8, 150 mM NaCl, 1mM EDTA). An excess of purified pr peptide was added to the virus (ratio 1:50) and the pH was changed with phosphate/citrate buffer to pH5.5-6-7-8. After 30 min incubation at 37°C the complex was pelleted in a SW55 rotor at 100 Kg for 1 hour and loaded on a 12% SDS gel. Antibody E21.3 was used in western blot to detect the viral E protein and antibody A3.2 was used to detect the pr protein. Band intensities were calculated in Image J software.

## ACKNOWLEDGEMENTS

We thank the staff at beamlines PX1 and PX2 at Soleil synchrotron (St. Aubin, France) and at PX beamlines at the ESRF (Grenoble, France); Franz X. Heinz and Karin Stiasny from Center for Virology, Medical University of Vienna, Austria for the gift of DENV pr protein and the plasmid for the production of ZIKV PF13 sE protein; Pablo Guardado-Calvo and Ignacio Fernandez from the Rey lab for helping with the MALS experiments and Alexander Rouvinski for help with the setting of the fusion experiments. This work was supported by the French ANR (Agence Nationale de la Recherche), grants ANR-17-CE15-0031-01 FLAVIMMUNITY to GBS.

