## Supplementary Material for "Molecular mechanisms regulating the pH-dependent pr/E interaction in yellow fever virus"

**SUPPL. FIGURE S1: Anion exchange purification of YFV pr/sE complex.**

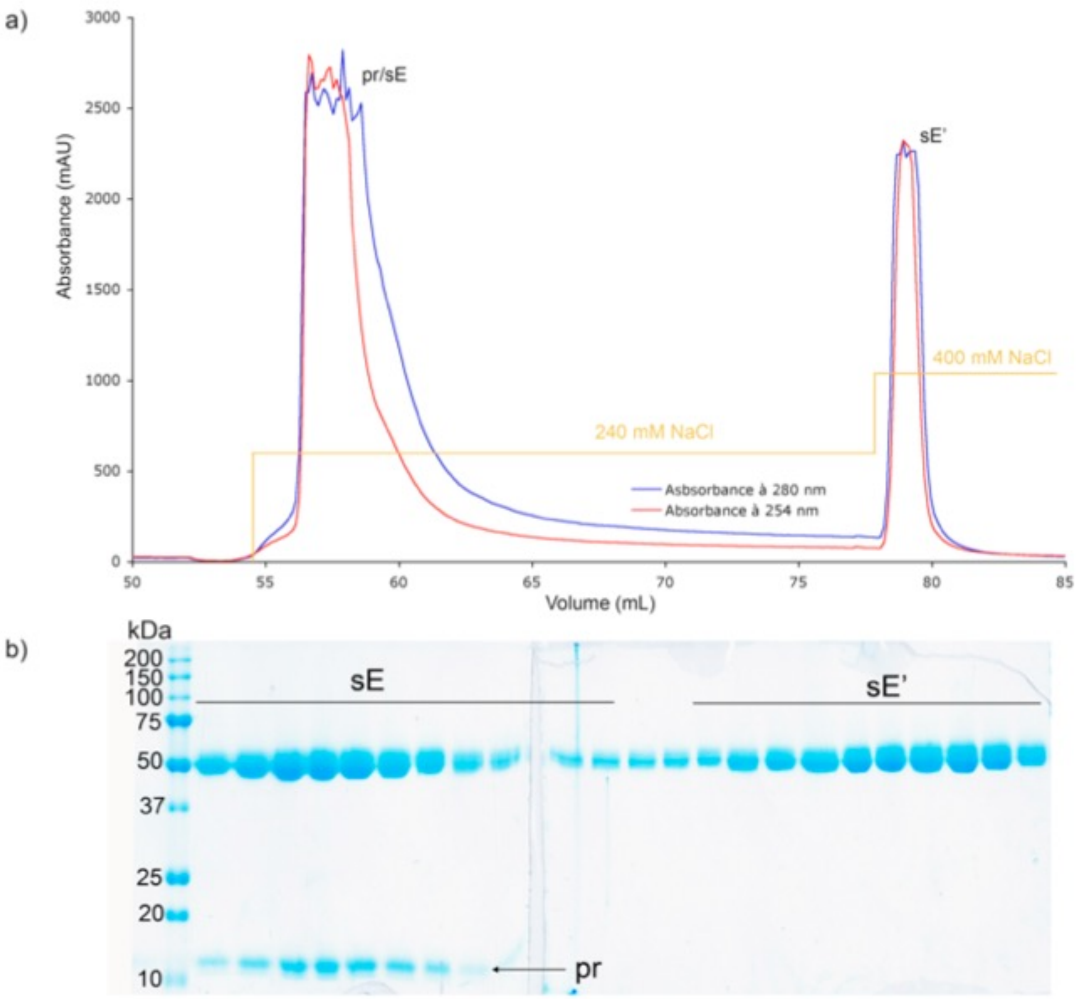


**Supplementary Figure S1 legend. (A)** SEC purified YFV pr/E protein was separated on a MonoQ 5/15 with a NaCl step gradient (240mM-400mM). **(B)** SDS-PAGE of fractions corresponding to the two peaks purified by Mono-Q column. Peak eluted at 240mM NaCl contains both sE and pr proteins. Peak eluted at 400mM NaCl contains only sE protein

**SUPPL. FIGURE S2: SEC-MALS elution volume profiles and SDS-PAGE of isolated sE and pr and of mixture of sE + pr at pH 8.0 and at pH 5.5 for ZIKV and DENV2 A259C mutant.**

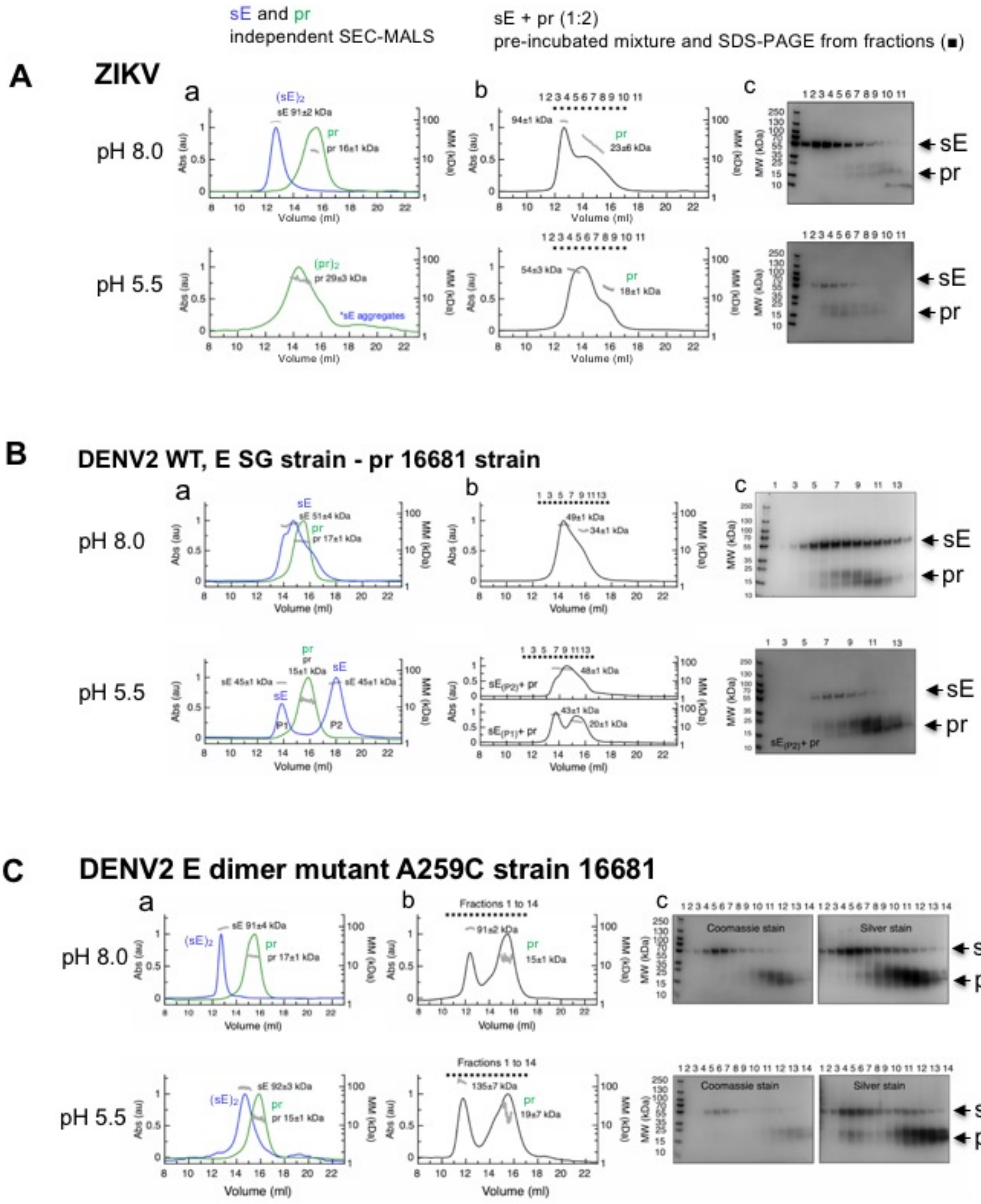

**Supplementary Figure S2 legend. (A, B,C)** For each virus, SEC-MALS elution volume profiles of sE, pr and pr:sE proteins and SDS-PAGE at two pH (pH 8, top panels; pH 5.5 bottom panels). **(a, b)** SEC-MALS profiles. Left y axis: the ultraviolet absorbance normalized by setting the highest peak to 1. Right y axis: molecular mass (kDa) determined by MALS, with the values for each species indicated on the corresponding peak. **(a)** SEC-MALS elution volume profiles of isolated sE (blue curves) and isolated pr (green curves) at pH 8.0 (top) and at pH 5.5 (bottom). **(b)** For each virus, SEC-MALS elution volume profiles of a mixture of sE with an excess of pr (1:2 sE:pr monomer:monomer molar ratio) equilibrated at pH 8.0 (top) and at pH 5.5 (bottom). The fractions analyzed by SDS-PAGE in (c) are indicated. **(c)** SDS-PAGE of the SEC-MALS fractions indicated in panels (b) at pH 8.0 (top) and at pH 5.5 (bottom). Coomassie blue staining. Only silver nitrate staining is shown for DENV2 sE A259C mutant at pH 8.0 (top) and at pH 5.5 (bottom).

**SUPPL. FIGURE S3: Quantification of E protein on YFV virus.**

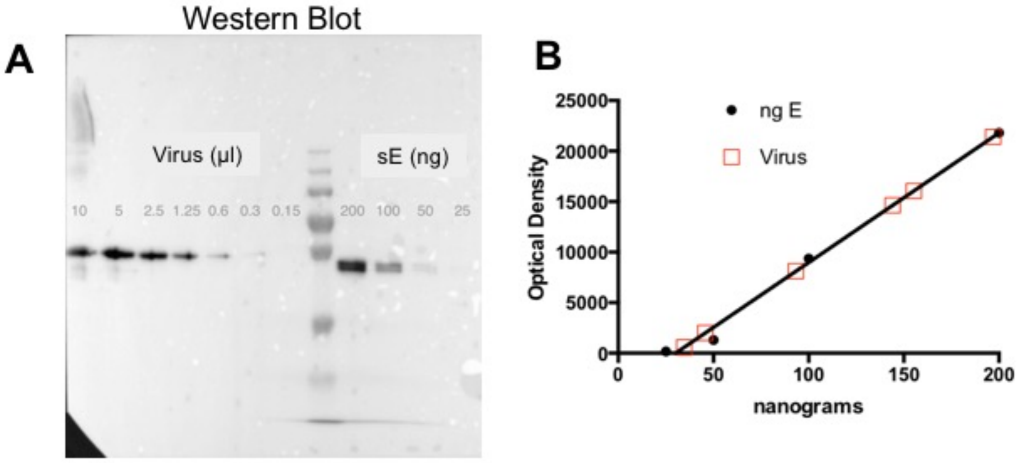


**Supplementary Figure S3 legend . (A)** Western-blot with an anti-E specific antibody of serial dilution of purified YF17D virus used for the fusion assays. The optical densities of the signal from the virus dilutions and known amounts of purified sE protein has been calculated using Image J software. **(B)** The amount of viral E protein has been then calculated by interpolation with the standard curve constructed with the values of the purified E protein using Prism 9.4 software.

### SUPPL. TABLE S1

**Supplementary Table S1. Data collection and refinement statistics.**

| | Asibi pr/sE | Asibi sE dimer |
|---|---|---|
| **PDB code** | 6EPK | / |
| ***Crystallization conditions*** | | |
| | 1.2-1.8M $Li_2SO_4$<br>0.1M Tris pH 8 | 1.26M $(NH_4)_2SO_4$<br>0.1M HEPES pH 7.5 |
| ***Data Collection*** [a] | | |
| Synchrotron | SOLEIL | SOLEIL |
| Beamline | Proxima-1 | Proxima-1 |
| Space group | $P\ 4_1$ | $P\ 4_3\ 2_1\ 2$ |
| Cell dimensions | | |
| a, b, c (Å) | 99.9, 99.9, 238.8 | 92.5, 92.5, 308.4 |
| α, β, γ (°) | 90, 90, 90 | 90, 90, 90 |
| Resolution (Å) | 30-2.7 (2.77-2.70) | 46.3-3.48 (3.81-3.48) [b] |
| R*merge* | 0.090 (1.123) | 0.011(2.20) |
| R*meas* | 0.111 (1.394) | 0.121 (2.28) |
| R*pim* | 0.063 (0.814) | 0.03 (0.60) |
| $< I / \sigma(I) >$ | 17.0 (1.2) | 13.1 (1.5) |
| $CC_{1/2}$ | 0.997 (0.519) | 0.999 (0.658) |
| Completeness (%) | 99.9 (100.0) | 62.8 (13.6) |
| *Ellipsoidal* [b] | | *93.4 (81.5)* |
| No. of Unique reflections | 63786 (4469) | 11358 (568) |
| Multiplicity | 5.4 (5.1) | 15.5 (13.8) |
| ***Structure determination*** | | |
| MR search models | 1SVB+3C5X | 6EPK |
| NCS | 2 | 2 |
| ***Refinement*** [a] | | |
| Resolution | 30.0-**2.7**<br>(2.75-2.70) | 30.0-**3.48**<br>(3.81-3.48) |
| No. of Work/Free reflections | 63742/3182<br>(2962/160) | 11613/983<br>(664/46) |
| Rwork/Rfree | 0.16/0.18<br>(0.34/0.37) | 0.25/0.27<br>(0.29/0.39) |
| Targeting | / | 6EPK |
| No. of atoms/waters | 7898/205 | 5964/0 |
| Rms deviations from ideality | | |
| Bond lengths (Å) | 0.003 | 0.006 |
| Bond angles (°) | 0.672 | 0.86 |
| Ramachandran Plot [c] | | |
| Favored (%) | 98.03 | 97.4 |
| Allowed (%) | 1.97 | 2.08 |
| Outliers (%) | 0.0 | 0.52 |
| Rotamers Outliers | 0.61 | 1.08 |

[a] Highest resolution shell is shown in parenthesis.
[b] Data collection statistics were computed after anisotropy correction by STARANISO.
[c] Ramachandran statistics were calculated with MolProbity.

PDB, Protein Data Bank; NCS, non-crystallographic symmetry; MR, molecular replacement.
Rms, root mean square; $CC_{1/2}$, correlation coefficient.
Rmeas, multiplicity-corrected R; Rpim, expected precision
